# Detecting Primary Progressive Aphasia (PPA) from Text: A Benchmarking Study

**DOI:** 10.1101/2025.02.19.639032

**Authors:** Ghofrane Merhbene, Fabian Lecron, Philippe Fortemps, Bradford C. Dickerson, Mascha Kurpicz-Briki, Neguine Rezaii

## Abstract

Classifying subtypes of primary progressive aphasia (PPA) from connected speech presents significant diagnostic challenges due to overlapping linguistic markers. This study benchmarks the performance of traditional machine learning models with various feature extraction techniques, transformer-based models, and large language models (LLMs) for PPA classification. Our results indicate that while transformer-based models and LLMs exceed chance-level performance in terms of balanced accuracy, traditional classifiers combined with contextual embeddings remain highly competitive. Notably, SVM using RoBERTa’s embeddings achieves the highest classification accuracy. These findings underscore the potential of machine learning in enhancing the automatic classification of PPA subtypes.

## 1 Introduction

Primary progressive aphasia (PPA) is a neurodegenerative disorder characterized by progressive language deficits as the primary symptom. It is typically classified into three subtypes (Gorno-Tempini et al., 2011): (1) the logopenic variant (lvPPA), associated with word-finding difficulties and impaired sentence repetition, often linked to Alzheimer’s pathology; (2) the semantic variant (svPPA), marked by deficits in word comprehension and object naming; and (3) the nonfluent variant (nfvPPA), characterized by effortful, halting, and telegraphic speech. The underlying pathology of svPPA and nfvPPA is often frontotemporal lobar degeneration (Rezaii et al., 2023). Diagnosing these subtypes traditionally requires extensive clinical assessment by expert neurologists, neuropsychologists, and speech-language pathologists, making the process resource-intensive and time-consuming. As a result, there is increasing interest in automated methods for efficient and accurate PPA classification. However, diagnosing PPA from textual data, such as transcripts of patient interviews, presents several challenges. The linguistic and syntactic markers that differentiate PPA subtypes are often subtle and overlapping, requiring robust feature extraction and classification techniques (Tippett, 2020). Furthermore, the limited availability of labeled clinical datasets and individual variability in language use exacerbate these challenges. Distinguishing svPPA from lvPPA is particularly difficult, as both subtypes involve word retrieval impairments. Despite these difficulties, accurate classification is crucial, given the distinct etiologies and treatment strategies associated with each PPA variant.

Recent advancements in natural language processing (NLP) have opened new avenues for automated diagnostic tools based on text. Prior research has demonstrated the potential of NLP in mental health assessment (Zhang et al., 2022), including applications in detecting bipolar disorder and schizophrenia (Aich et al., 2022). However, research on applying NLP to neurodegenerative diseases, particularly PPA, remains limited. Notably, there is a lack of systematic benchmarking studies that compare multiple computational approaches for PPA classification. To address this gap, we conduct a comprehensive benchmarking study, systematically evaluating a diverse range of models, from traditional machine learning (ML) methods with various feature extraction techniques to transformer-based models and large language models (LLMs). By providing a comparative analysis of these approaches, our study offers new insights into the effectiveness of different computational techniques for the automated classification of PPA subtypes.

## 2 Related Work

Research on PPA has primarily focused on understanding its clinical subtypes and linguistic manifestations (Henry et al., 2016). Studies in clinical neurology and neuropsychology have detailed the unique language impairments associated with lvPPA, svPPA, and nfvPPA, highlighting the importance of linguistic and syntactic analysis in diagnosis (Wauters et al., 2023). However, leveraging computational methods for the diagnosis of PPA remains an emerging area.

In the field of natural language processing (NLP), traditional machine learning models have been widely applied to clinical text classification tasks, including disease detection and subtype identification. In their study, Fraser et al. (2014) explored the use of computational linguistics for identifying different variants of PPA. They compared various feature sets, including acoustic, lexical, and syntactic features, and demonstrated that combining multiple modalities significantly improved classification performance. Their findings highlighted the importance of leveraging diverse linguistic markers to distinguish PPA subtypes, particularly the nonfluent variant (nfvPPA), which often exhibits clear syntactic deficits. Similarly, Themistocleous et al. (2021) achieved a classification accuracy of 80% by combining acoustic and linguistic features and using them as input for a deep neural network model. Building on this foundation, Rezaii et al. (2022) investigated the relationship between lexical and syntactic complexity during language production in individuals with PPA and healthy controls. Their study identified a syntax-lexicon trade-off where individuals with syntactic deficits, such as those with nfvPPA, used semantically richer words, while those with lexicosemantic deficits (e.g., svPPA or lvPPA) produced syntactically complex sentences. Their approach achieved a classification accuracy of up to 92% when distinguishing nfvPPA in a one-vs-all setup. In more recent work, Rezaii et al. (2024) explored the use of large language models (LLMs) to classify PPA subtypes based on connected speech. Their approach incorporated verb frequency and other linguistic features to align text-based speech patterns with brain scan findings, achieving 88.5% agreement on PPA clusters with LLMs. A supervised classifier using features identified by the LLM further improved accuracy to 97.9%. This study highlights the potential of LLMs in identifying linguistic markers of PPA subtypes and represents a significant advance in the application of NLP to clinical tasks. Cong et al. (2024b) also investigated the use of LLMs for detecting the presence, subtypes, and severity of aphasia in both English and Mandarin Chinese speakers. Their findings revealed that applying LLMs without finetuning resulted in accuracy levels close to chance for aphasia subtyping.

Language impairments, such as PPA, are often among the earliest signs of broader cognitive decline, including dementia (Harvard Health Publishing, 2022). Santander-Cruz et al. (2022) employed a combination of syntactic and semantic analyses to detect dementia in transcribed data from the Pitt Corpus database provided by DementiaBank^1^. They extracted features such as spelling mistakes, grammar errors, and cosine similarity and evaluated their effectiveness using machine learning models, including SVMs and neural networks. Notably, syntactic features alone achieved an F1-score of 77% with SVMs. While their approach demonstrated the effectiveness of syntactic features, it remained limited in scope, focusing on a predefined feature set and a small selection of models. In contrast, our study systematically evaluates a wider range of methodologies, from traditional machine learning models with different feature extraction techniques to transformer-based models and LLMs, to comprehensively assess the potential of NLP techniques for PPA classification.

## 3 Dataset

### 3.1 Overview

The data used in this study was shared with us by (Rezaii et al., 2022). The dataset consists of clinical transcripts from interviews with individuals diagnosed with one of the PPA subtypes, as well as control participants without a PPA diagnosis. All participants were shown a drawing of a family at a picnic from the Western Aphasia Battery-Revised (Clark et al., 2020) and were asked to describe it using as many full sentences as possible. A total of 79 interviews with PPA patients were sourced from a study conducted within the PPA program at the Frontotemporal Disorders Unit of Massachusetts General Hospital (MGH). Expert neuropsychiatrists and speech-language pathologists carried out the assessment and annotation. The dataset also includes 53 healthy controls, sourced from the Speech and Feeding Disorders Laboratory at Massachusetts General Hospital (MGH) and Amazon’s Mechanical Turk (MTurk). The distribution of subtypes is shown in Table 1. All participants were native English speakers with no self-reported history of brain injury or speech/language disorders. Healthy controls and PPA patients were matched in terms of age, gender, handedness, and years of education.

**Table 1:**
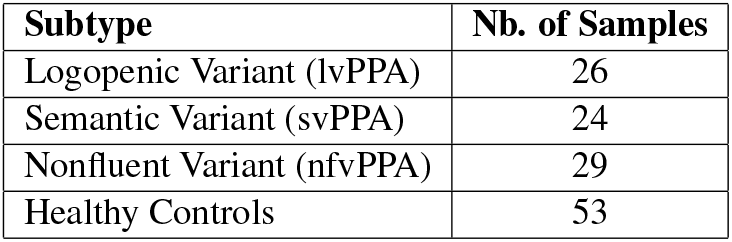
Distribution of subtypes and number of samples in the original version of the dataset.

Two versions of the dataset were used throughout this study:

- **Original version:** As described in the paragraph above.
- **Expanded version:** For each patient/participant, the transcript was split sentence-wise, with each sentence labeled identically to the original full transcript. This resulted in the label distribution provided in Table 2.

**Table 2:** Distribution of subtypes and number of samples in the expanded version dataset.

| Subtype | Nb. of Samples |
| --- | --- |
| Logopenic Variant (lvPPA) | 433 |
| Semantic Variant (svPPA) | 402 |
| Nonfluent Variant (nfvPPA) | 335 |
| Healthy Controls | 960 |

### 3.2 Data Preprocessing

Preprocessing the data is a crucial step before applying machine learning models, as it ensures the integrity of the linguistic and syntactic features. This section details the preprocessing steps undertaken to prepare the dataset.

The first step included converting all text to lowercase to standardize case sensitivity. Special characters were removed, retaining only intentionally alphanumeric characters and punctuation marks, as these features are significant in the diagnosis of PPA. For instance, punctuation patterns can signify pauses, sentence boundaries, or telegraphic speech, which are critical markers for distinguishing between PPA subtypes. Nonfluent Variant PPA (nfvPPA), in particular, is marked by halting speech and frequent pauses. Following this, the text was tokenized into individual words for further analysis.

It is important to mention that the preprocessing steps were performed for the experiments with the traditional machine learning (ML) models described in Section 4.2.

## 4 Methodology

### 4.1 Evaluation Reference Points

To evaluate the performance of the models in this multi-class classification task, we define two reference metrics to provide a point of comparison for balanced accuracy:

- **Naïve Random Reference (chance level):** Assuming a random guess, each of the four classes has an equal probability of being selected, resulting in a uniform class probability of 25%.
- **Stratified (Weighted) Random Reference:** This reference metric accounts for class imbalance by weighting each class proportionally to its frequency in the dataset. Since this metric incorporates dataset imbalance, it provides a more realistic reference than uniform random guessing.

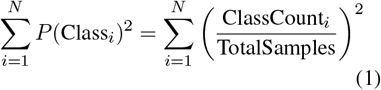

### 4.2 Traditional Machine Learning Models

The initial experiments in this benchmarking study involve applying various feature extraction techniques in combination with a predefined set of traditional machine learning models. The following subsection provides an overview of the feature extraction techniques used.

#### 4.2.1 Feature Extraction techniques

Several feature extraction strategies were used in this investigation. Traditional techniques, such as **TF-IDF** (Salton and Buckley, 1988) and **Bag of Words** (BoW) (Harris, 1954), emphasized word frequency and document-specific relevance. Word embeddings like **Word2Vec** (Mikolov et al., 2013), **FastText** (Bojanowski et al., 2017), and **GloVe** (Pennington et al., 2014) capture semantic relationships in dense vector spaces. **FastText** also uses subword information. Transformer-based models, such as **BERT** (Devlin et al., 2019) and **RoBERTa** (Liu et al., 2019), were used to produce contextual embeddings, resulting in a semantic understanding. **N-grams** (bigrams and 4-grams) (Brown et al., 1992) captured local context, whereas **LSA** (Deerwester et al., 1990) and **LDA** (Blei et al., 2003) detected latent structures and topics. **Dependency parsing** (Kiperwasser and Goldberg, 2016) examined grammatical links to extract syntactic elements important for diagnosing language problems.

#### 4.2.2 Machine Learning (ML) Models

Traditional ML models have helped advance artificial intelligence and continue to offer advantages such as interpretability, computational efficiency, and adaptation to smaller datasets (Murphy, 2012). Despite the growing dominance of large language models (LLMs), the performance of traditional models should not be overlooked, particularly in tasks where linguistic and syntactic features play a central role.

To ensure a robust benchmarking process, we incorporate five widely-used traditional machine learning models: Support Vector Machine (SVM), Naive Bayes (NB), Logistic Regression (LR), Multilayer Perceptron (MLP), and XGBoost. These models were evaluated in combination with the feature extraction techniques detailed in the previous section.

Considering the limited sample size, default parameter settings without hyperparameter finetuning were used for all models, ensuring simplicity and reproducibility in the benchmarking process. Experiments were conducted using stratified 5-fold cross-validation to mitigate overfitting and provide reliable performance estimates. In total, 55 experiments were only performed (5 classifiers × 11 feature extraction techniques).

### 4.3 Transformer-based Models

In addition to traditional machine learning models, this study evaluates the performance of transformer-based models, which have revolutionized natural language processing by taking advantage of attention mechanisms and contextual embeddings. These models are particularly well-suited for tasks involving subtle syntactic variations and capturing long-term dependencies, making them strong candidates for the classification task at hand. While some transformer models were previously used to generate embeddings for feature-based approaches (as detailed above), here, they are directly employed as classifiers to assess their full predictive capabilities.

The transformer-based models included in this benchmarking study are as follows:

- **BERT**: A bidirectional transformer that captures context from both left and right of a word, making it effective for tasks that require deep semantic understanding (Devlin et al., 2019).
- **RoBERTa**: A robustly optimized version of BERT with improved training strategies and increased training data, designed to improve performance on a variety of NLP tasks (Liu et al., 2019).
- **MentalBERT**: A domain-specific transformer model fine-tuned on mental health-related text, aimed at capturing linguistic patterns specific to this domain (Ji et al., 2022).
- **ClinicalBERT**: A transformer fine-tuned on clinical text, optimized for healthcare-related tasks and well-suited for datasets with medical or diagnostic data (Alsentzer et al., 2019).

These models are evaluated using the same crossvalidation protocols applied to traditional ML models, ensuring fair comparison.

## 5 Large Language Models (LLMs)

Large Language Models (LLMs) represent a significant breakthrough in artificial intelligence, demonstrating exceptional capabilities across a wide range of NLP tasks. These models, like OpenAI’s GPT series and Google’s Gemini, are built upon transformer-based architectures and are known by their immense size, comprising billions or even trillions of parameters. Their extensive training, combined with their parameterization, allows them to achieve high performance in a wide range of NLP tasks, including text generation.

In this study, we employ a prompt-based approach to leverage LLMs for the classification of PPA subtypes. Rather than fine-tuning these models, we evaluate their few-shot performance by designing a structured prompt tailored to our classification task. The following large language models (LLMs) were used in this study:

- **LLAMA:** meta-llama/ Meta-Llama-3-8B-Instruct, sourced from the Hugging Face repository, developed by Meta, with 8 billion parameters, fine-tuned for instruction-based tasks (Touvron et al., 2023).
- **Mistral:** mistralai/Mistral-7B-Instruct-v0.2, sourced from the Hugging Face repository, developed by Mistral AI, with 7 billion parameters, optimized for instruction-based and conversational tasks (Jiang et al., 2023).
- **GPT-3.5-turbo:** Developed by OpenAI, a 175 billion parameter model known for its general-purpose conversational and reasoning capabilities (Brown et al., 2020).
- **GPT-4o-mini:** Developed by OpenAI, a lightweight variant of GPT-4, fine-tuned for optimized performance on smaller computational setups (OpenAI, 2023).

The original version of the data was used, and the prompt was carefully designed in collaboration with a clinical expert in the field (see Appendix A).

The temperatures used for each model are presented in Table 3. For Mistral and LLAMA, we used a relatively low temperature (0.2) to ensure more deterministic outputs^2^, given that these models can occasionally exhibit excessive variation at higher temperatures. In contrast, GPT-3.5 and GPT4o-mini were assigned a moderately higher temperature (0.7) to encourage more diverse responses while maintaining overall coherence.

**Table 3:**
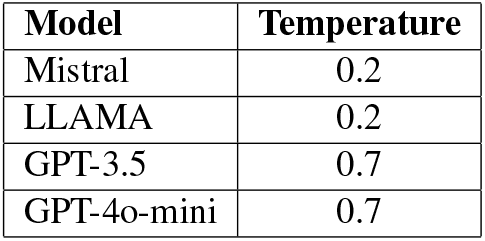
Temperature used for each model.

## 6 Results

To ensure a comprehensive evaluation, we rely on widely recognized classification metrics, including balanced accuracy, weighted F1-score, weighted precision, weighted recall, Area Under the Curve (AUC), as well as a confusion matrix for LLM based experiments. All experiments are evaluated using 5-fold cross-validation to ensure robustness and minimize overfitting. The results are presented as bar charts, with balanced accuracy’s reference performance indicated by vertical lines to provide a clear point of comparison.

### 6.1 Traditional Machine Learning Models

Figures 1 and 2 present the performance of the top-performing traditional machine learning models (in terms of F1-score), namely SVM and LR. Each colored bar represents the machine learning model paired with a different feature extraction technique. The results for the other models, including MLP, Naïve Bayes, and XGBoost, are provided in Appendix C for completeness.

**Figure 1:**
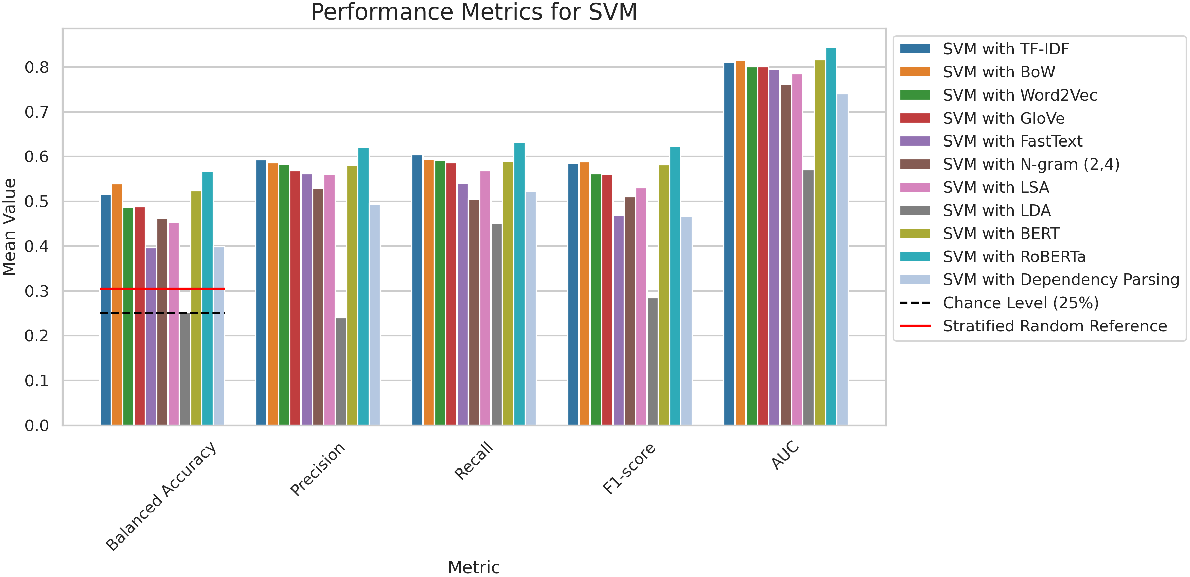
SVM performance.

**Figure 2:**
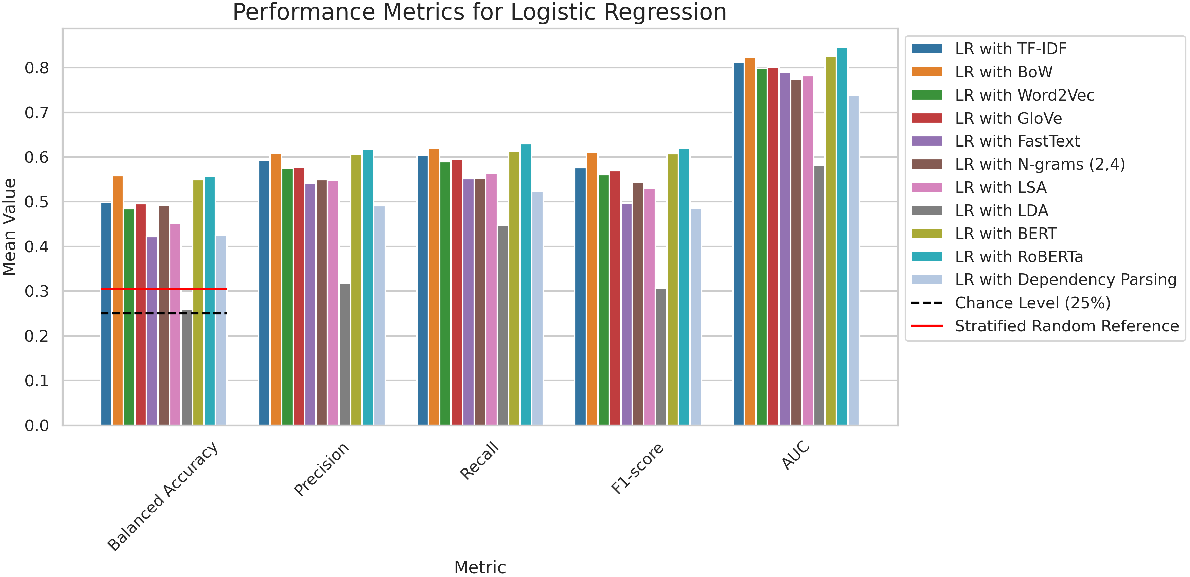
Logistic Regression performance.

In terms of balanced accuracy, using RoBERTa’s features yielded the best results across almost all models, while LR showed a tie between the use of RoBERTa and the Bag-of-Words features. RoBERTa, followed closely by BERT, also demonstrated superior performance across other metrics, including weighted precision, weighted recall, weighted F1-score, and AUC. Specifically, RoBERTa achieved over 60% weighted precision, weighted recall, and F1-score for both SVM and LR, while exceeding 80% AUC for SVM, MLP, LR, and XGBoost.

### 6.2 Transformer-based Models

Figure 3 illustrates the performance of the various transformer-based classifiers. All models significantly outperform the reference metrics in terms of balanced accuracy, with RoBERTa and BERT demonstrating comparable top-tier performance, closely followed by MentalBERT. Regarding the F1-score, RoBERTa and BERT achieve the highest results, both exceeding 60%, with RoBERTa exhibiting a slight advantage. Similar trends are observed for weighted precision and weighted recall, where RoBERTa, BERT, and MentalBERT all achieve scores over 60%. In terms of AUC, RoBERTa, MentalBERT, and BERT all demonstrate strong performance, achieving results above 80%.

**Figure 3:**
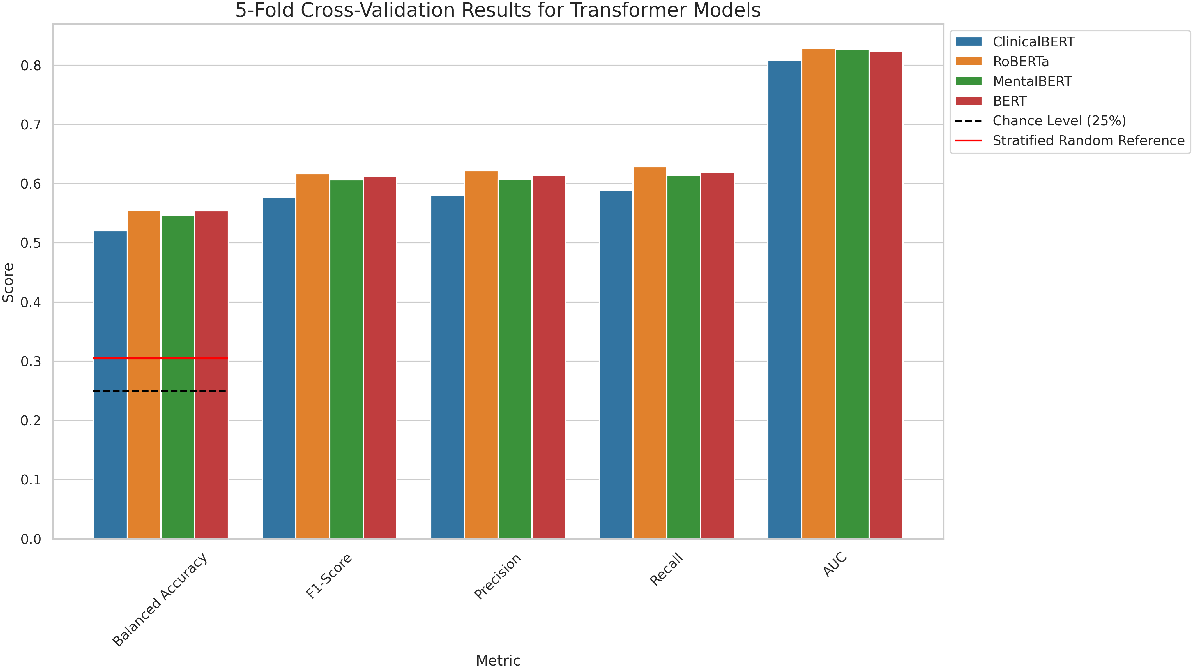
Transformer-based models performance.

### 6.3 Large Language Models (LLMs)

Figure 4 presents a bar chart illustrating the performance of LLAMA, which achieved the highest balanced accuracy, weighted precision, weighted recall, and F1-score among all models. For completeness, the results of Mistral, GPT-3.5-turbo and GPT-4o-mini are provided in Appendix D. All models, except for Mistral, outperformed our reference metrics, with LLAMA achieving a balanced accuracy of 63%.

**Figure 4:**
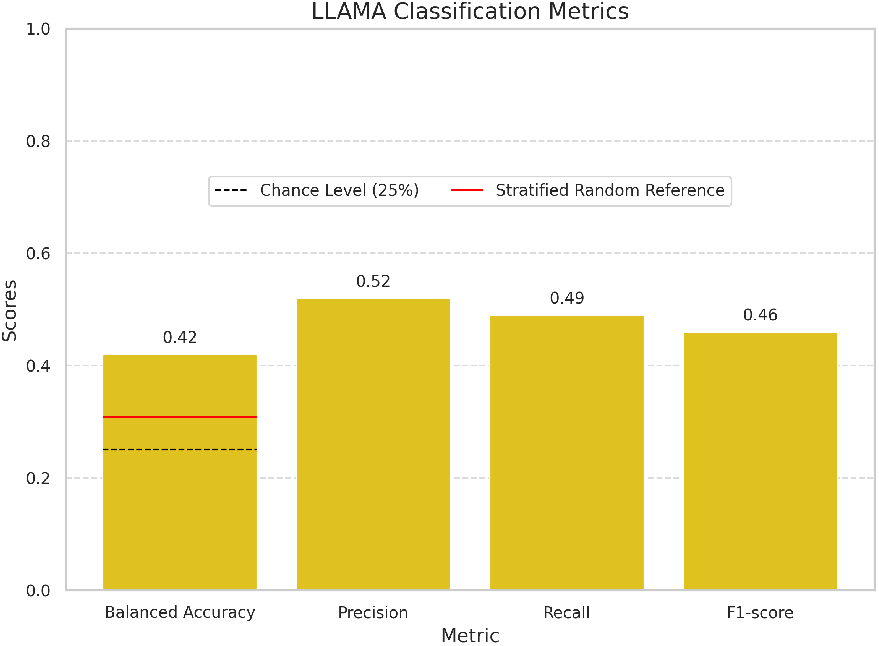
LLAMA Performance.

**Figure 5:**
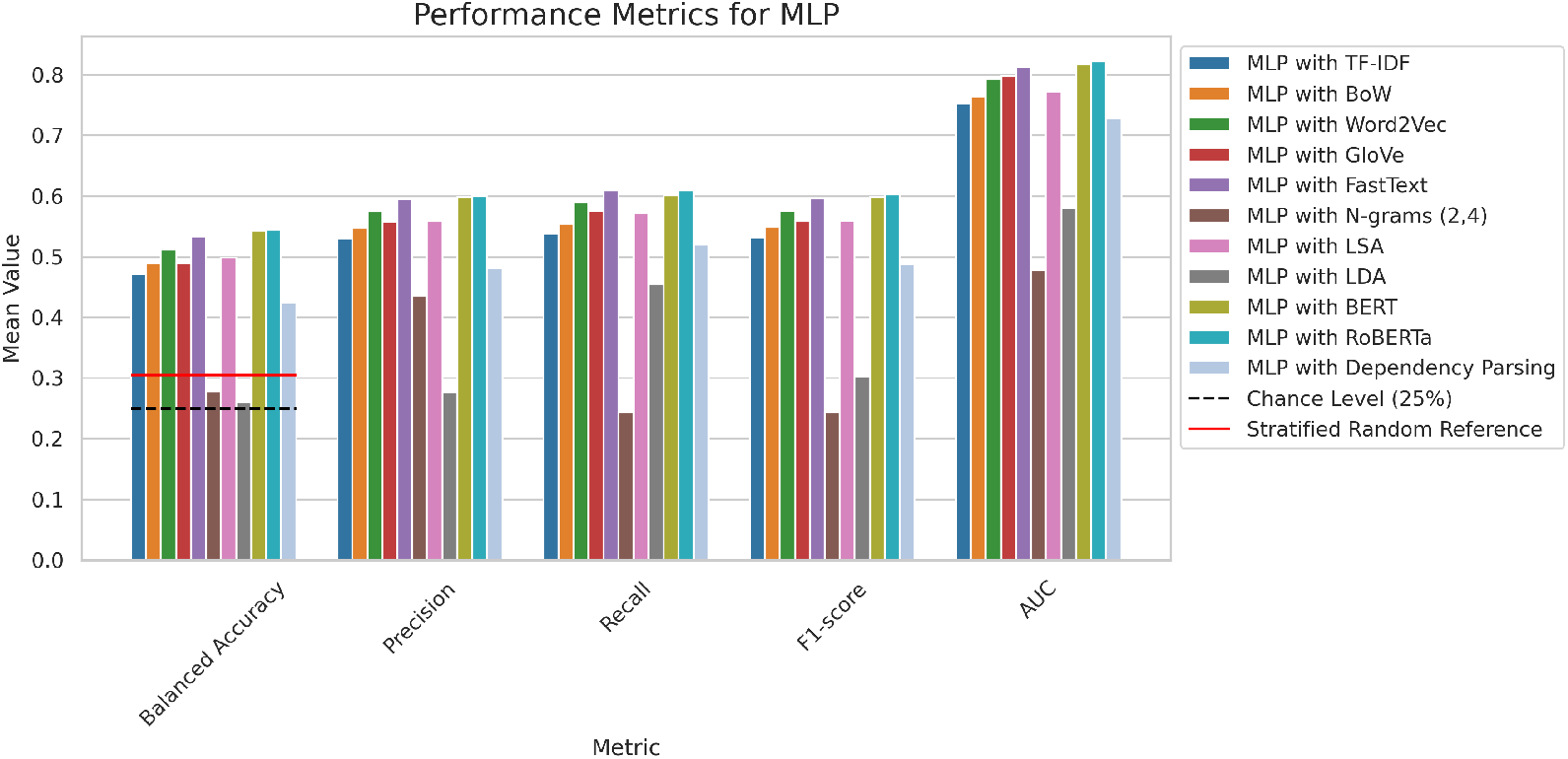
MLP performance.

**Figure 6:**
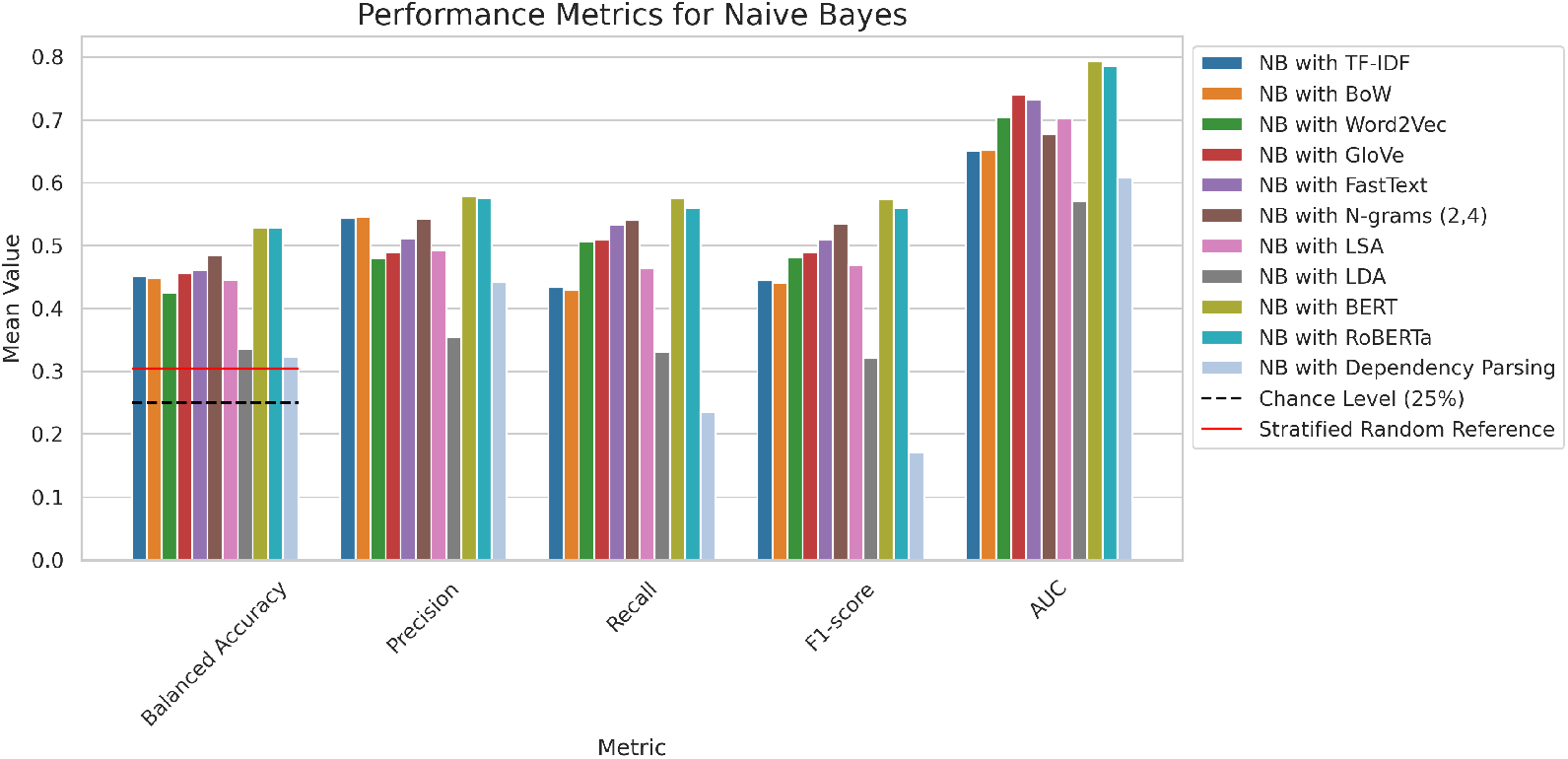
Naive Bayes performance.

**Figure 7:**
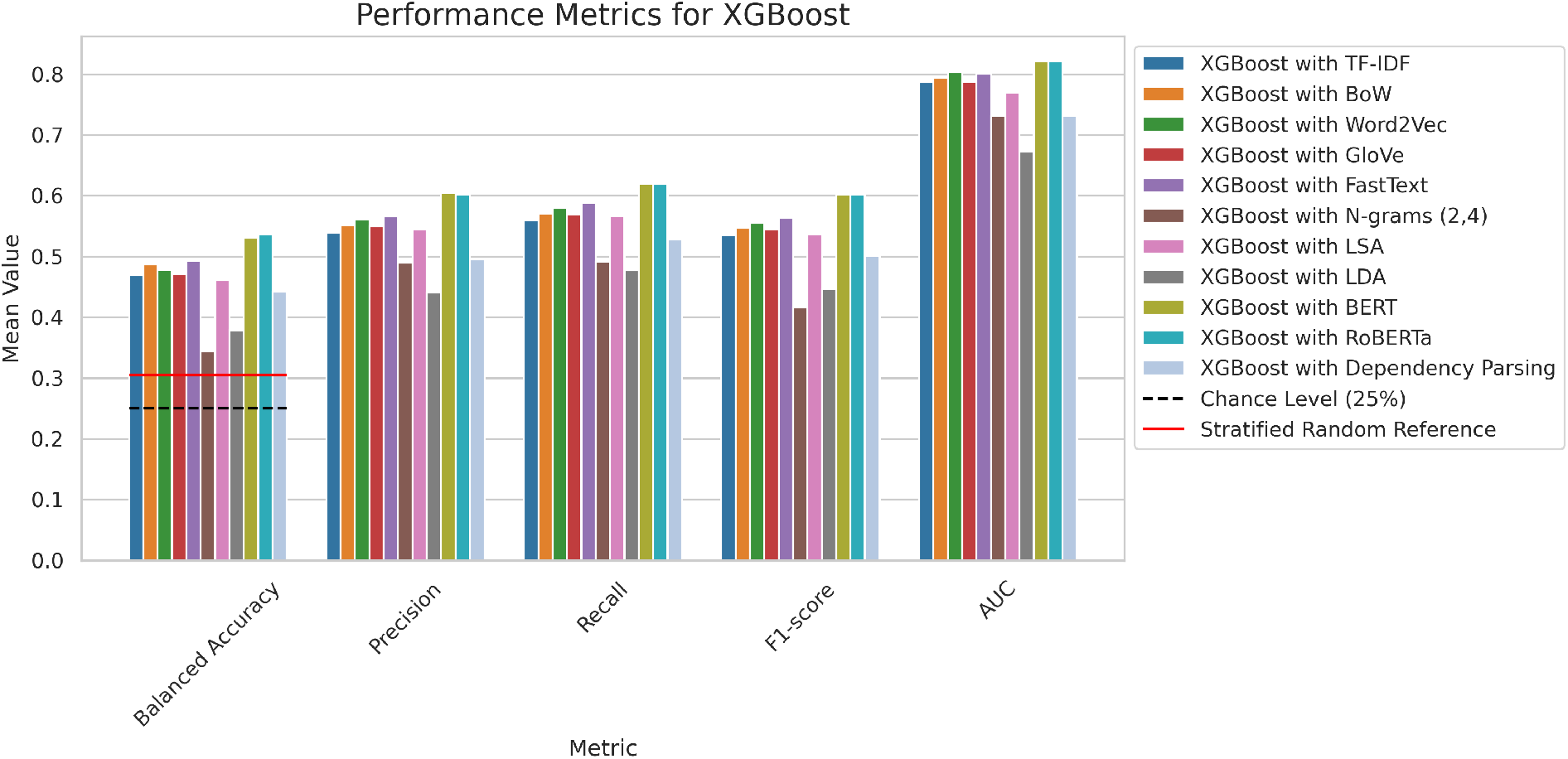
XGBoost performance.

**Figure 8:**
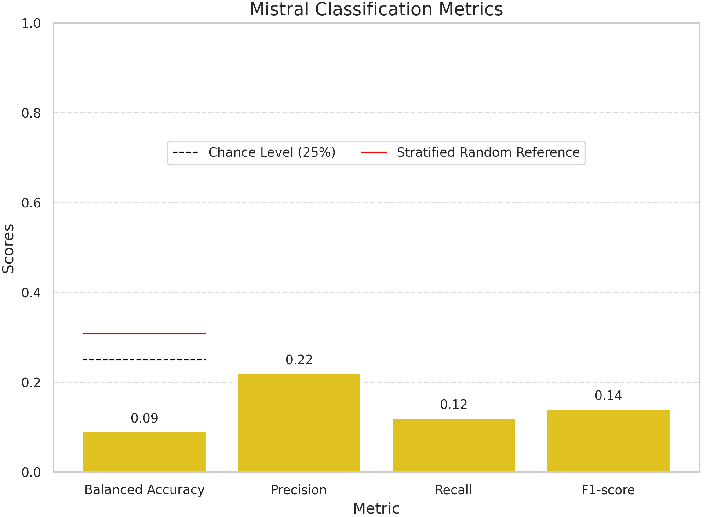
Mistral Performance.

**Figure 9:**
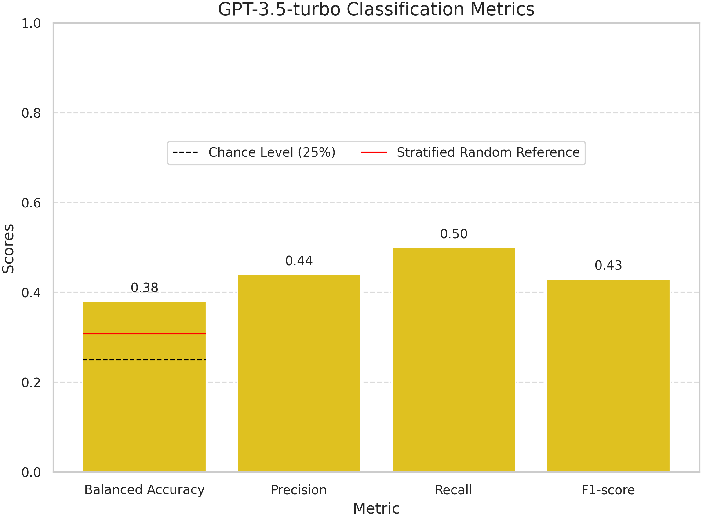
GPT-3.5 Performance.

**Figure 10:**
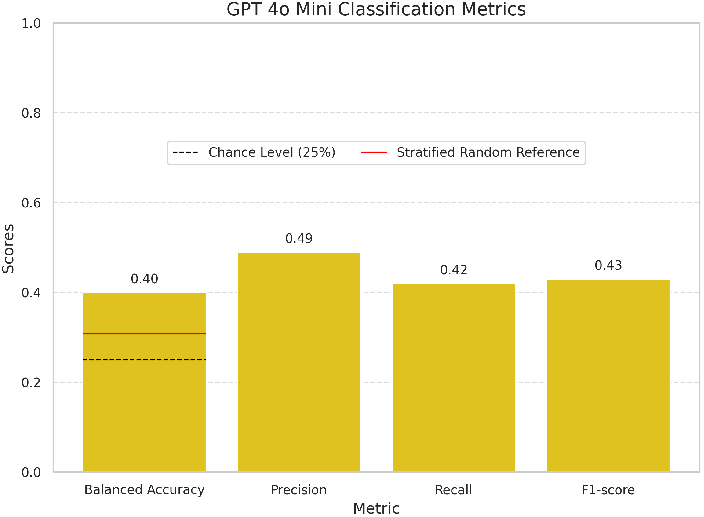
GPT-4o-minia performance.

Figures 11, 12, 13, and 14 in Appendix D present the confusion matrices for all four models. For LLAMA, we observe a strong performance in correctly predicting both the control group and lvPPA, but the model struggled with predicting any svPPA samples. Mistral’s performance, shown in Figure 12, was the weakest, as it assigned multiple times the label, *unknown*, when it failed to classify a sample correctly. Both GPT models performed well in identifying the control group, with GPT-3.5Turbo showing a slight edge over the other model. However, both models faced significant difficulty with lvPPA. GPT-3.5 also showed limited success with svPPA, whereas GPT-4o-mini performed better on both svPPA and nfvPPA.

**Figure 11:**
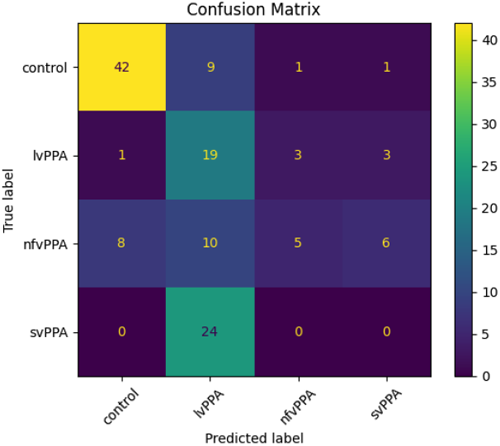
LLAMA Confusion Matrix.

**Figure 12:**
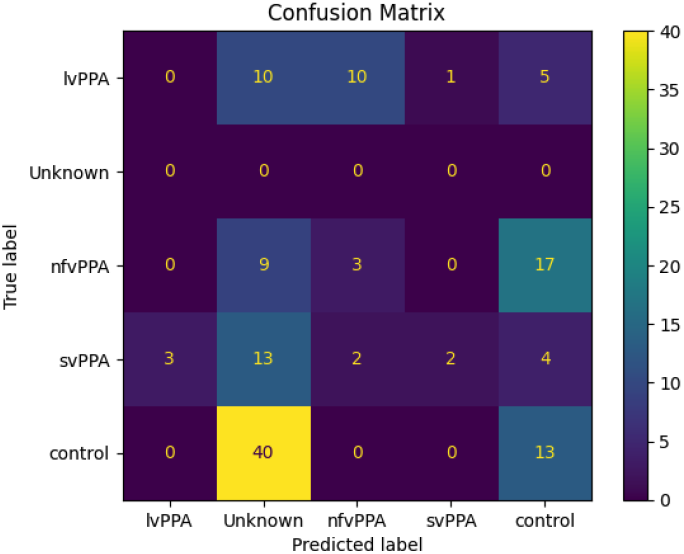
Mistral Confusion Matrix.

**Figure 13:**
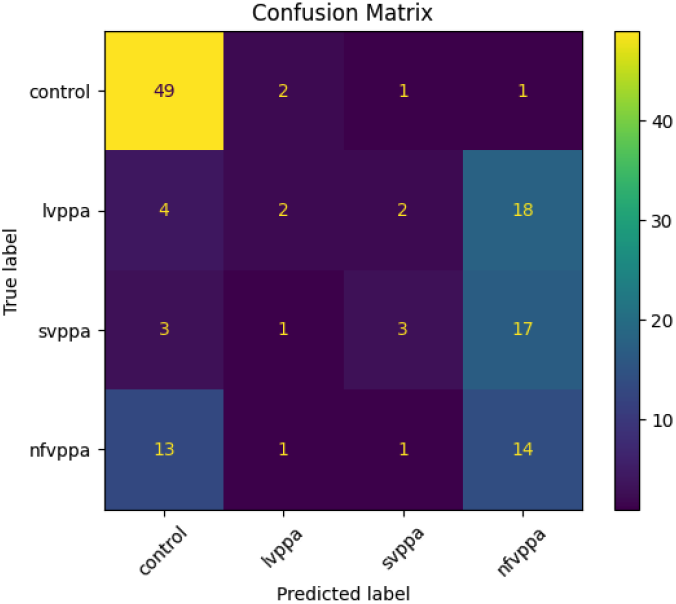
GPT-3.5-Turbo Confusion Matrix.

**Figure 14:**
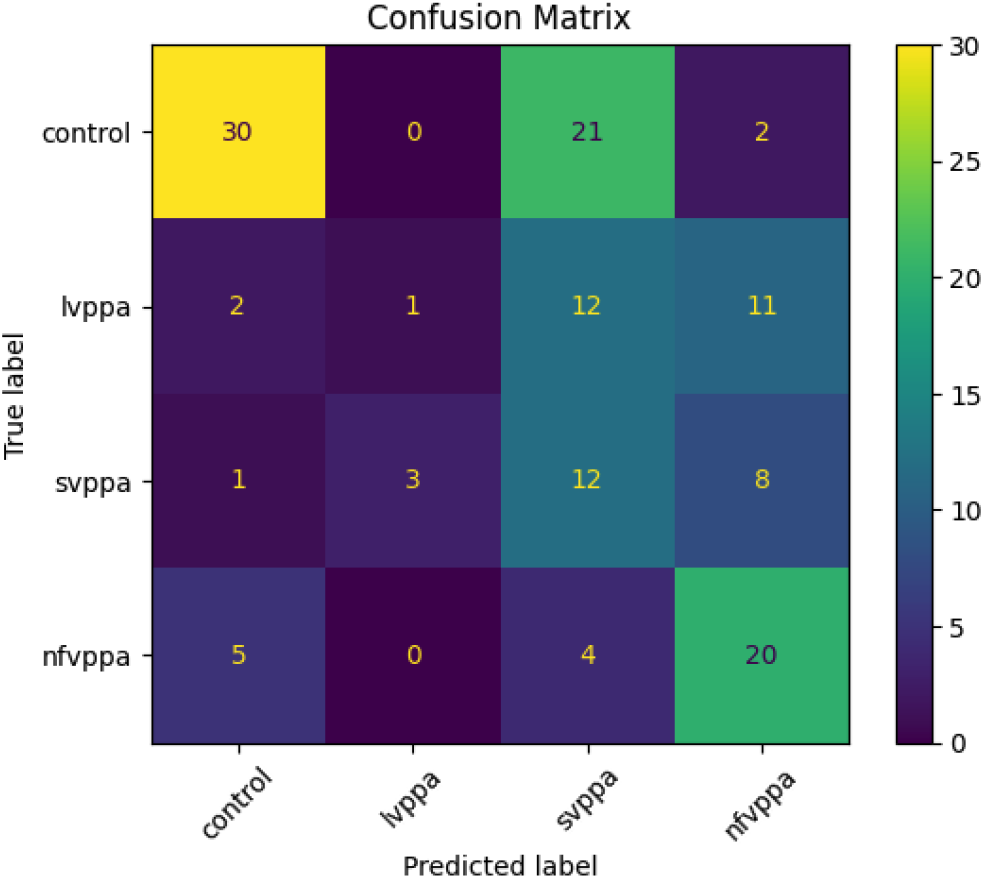
GPT-4o-mini Confusion Matrix.

## 7 Discussion

The results of this study provide valuable insights into the potential of various models for detecting primary progressive aphasia (PPA) subtypes. Benchmarking traditional machine learning approaches, transformer-based models, and large language models (LLMs) holds significant importance in advancing clinical diagnostics. These efforts not only reveal key trends and performance disparities but also underscore the broader potential of these models to improve the detection and classification of complex clinical conditions, such as PPA.

The results from traditional machine learning models reveal the critical role of feature extraction in determining performance. Across most models, RoBERTa’s embeddings consistently yielded superior results compared to traditional feature engineering techniques. This was particularly evident in SVM and LR, where the use of RoBERTa features resulted in over 60% weighted precision, weighted recall, and F1-scores, as well as AUC values exceeding 80%. These findings highlight the potential of combining robust feature extraction methods with simpler classifiers to achieve competitive results, especially in resource-constrained environments. In addition, in use cases where context is important relying on contextual embeddings like those generated by RoBERTa is generally expected to yield better results. The LR model’s strong performance with BoW further suggests that simpler techniques may still be viable in certain scenarios, particularly when interpretability is prioritized (Itani et al., 2019). When dealing with sensitive medical conditions such as PPA, interpretation is paramount, as clinicians and researchers need to understand the rationale behind model predictions. The ability to explain why a model classified a patient’s condition can thus foster trust.

In line with recent work by Rezaii et al. (2022), our findings further emphasize the inherent difficulty of the multi-class classification task for PPA subtypes. Relying on a syntax-lexicon approach, the authors achieved a high accuracy (92%) in a binary classification task but reported a significant drop to 66% accuracy in multi-class classification. This stark contrast underscores the challenges faced by the overlapping symptoms and complexity of different PPA subtypes. Similar to their findings, our results confirm that advanced machine learning techniques, while promising, still face limitations when addressing multi-class classification in this domain.

Furthermore, transformer-based models such as RoBERTa achieved balanced accuracy and F1scores exceeding 60%, which, highlights the intrinsic challenges of capturing the subtle linguistic and syntactic variations inherent in PPA subtypes especially in a multi-class classification setting. These results align with the broader challenges outlined by Gorno-Tempini et al. (2011), who discuss the diagnostic complexity of PPA due to the heterogeneity and overlapping symptoms among its subtypes. The strong performance of transformers in our study reaffirms their potential to address these challenges, though further refinement is necessary to improve accuracy. This finding was also emphasized by Cong et al. (2024a), where the authors reaffirmed the potential of transformer-based models in healthcare, particularly in identifying complex patterns essential for the early detection and classification of neurodegenerative diseases. In addition, an important insight is that general-domain models appear to outperform domain-specific ones. Specifically, RoBERTa and BERT consistently produced stronger results than ClinicalBERT and MentalBERT, although MentalBERT’s performance remained reasonably close on most metrics. One possible explanation is that larger, more diverse pretraining corpora may help general-domain models capture a wider range of linguistic cues. However, even if domain-specific models are adjusted to specialised vocabulary, they could overlook some contextual cues or universal language patterns that are useful in broader tasks. In fact, general-domain BERT can occasionally stay competitive or even outperform specialised models, according to Alsentzer et al. (2019), indicating that in some situations, greater coverage may outweigh niche specialization in certain scenarios. While LLMs outperformed our reference metrics in terms of balanced accuracy (with the exception of Mistral), their results were inconsistent across the subtypes (see confusion matrices in Appendix D). LLAMA achieved the highest balanced accuracy, weighted precision, weighted recall, and F1-score, yet it struggled particularly with svPPA classification. One likely explanation is that we used these models without fine-tuning, relying solely on prompting. Unlike smaller models explicitly optimized for classification through feature-based learning, LLMs generate responses based on broad language modeling objectives, which may not align well with structured clinical classification. These results highlight the limitations of few-shot LLM classification, where performance may be constrained without fine-tuning or domain adaptation.

Table 4 highlights the best-performing models across our experiments. While most models demonstrated comparable performance, LLAMA stood out negatively; despite outperforming other LLMs, it failed to match the top models in other categories. SVM paired with RoBERTa embeddings emerged as the strongest model, achieving the highest scores in balanced accuracy, weighted F1-score, weighted precision, and weighted recall, though by a narrow margin.

**Table 4:**
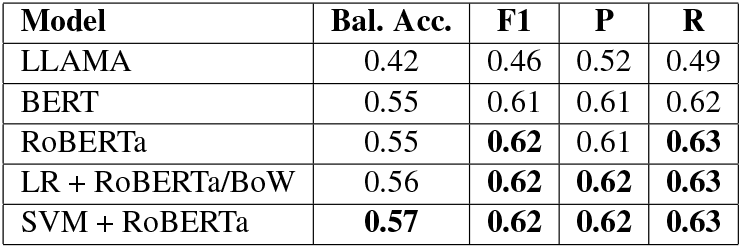
Performance metrics for the best models (Bal. Acc. = Balanced Accuracy, P = Precision, R = Recall). Highest value(s) in each column are in bold.

## 8 Conclusion

Our findings show the promise of using AI in the classification of PPA subtypes. The results demonstrate that although transformer-based methods sometimes yield slightly higher metrics, they do not decisively outperform classical feature based techniques such as SVM paired with RoBERTa’s embeddings. This highlights the inherent complexity of the classification task, shaped by the overlapping symptoms across PPA subtypes. Given the limitations observed in prompt-based LLM experiments, future work should explore task-specific fine-tuning to better align these models with the linguistic characteristics of PPA. Further error analysis may also provide insights into systematic misclassifications, guiding refinements in model training.

## 9 Limitations

The task of classifying primary progressive aphasia (PPA) subtypes presents a significant challenge due to the overlapping symptoms and linguistic impairments between subtypes. Additionally, our dataset while useful for benchmarking remains relatively small and lacks demographic metadata, preventing an analysis of potential biases across different population groups. Computational constraints also limited our ability to explore hyperparameter tuning for all our experimentss, which may have impacted model performance. This is particularly relevant for traditional classifiers and transformer-based models, where optimal settings could have led to improved results. Similarly, our exclusive reliance on natural language prompts for LLMs (although designed with expert input) may have limited their performance, as we lacked fine-tuning or deeper insights into their decision-making processes. Without explicit control over how LLMs generate classifications, their outputs can be difficult to interpret and optimize for this task. Future work should explore fine-tuning approaches and systematic hyperparameter optimization to better align model performance with the complexities of PPA classification.

## 10 Ethical Considerations

The dataset used in this research was anonymized and sourced from a previously published study. This ensures that the privacy and data protection of the original participants are upheld. However, due to the anonymization process, we have limited information about participants’ demographic backgrounds. As a result, we cannot assess potential biases or limitations of our classifiers across different societal groups. To ensure broader applicability and fairness, it is essential to validate our findings on a larger and more diverse dataset before deploying such a system in real-world scenarios.

Additionally, while this work does not directly create an automated diagnostic tool, its findings could contribute to the development of such technologies in the future. We emphasize that the goal is to assist clinicians rather than replace them, and we acknowledge the potential risk of misuse if such tools were to be used as substitutes for expert judgment.

## A Prompt for Clinical Text Classification

The following prompt was used to guide the clinical text classification task performed by the large language models:

You are a clinical text classifier specializing in language and speech characteristics related to Primary Progressive Aphasia (PPA). Based on the provided interview transcript of a patient, classify the text into one of the following categories:

- **lvPPA**: Logopenic Variant, Characterized by word-finding difficulties and impaired repetition abilities. Patients may frequently pause or hesitate as they search for words, and they may struggle to repeat phrases accurately. Example: Patient might say, “I went to the… um… place where… you know, people get… books,” when trying to say “library.” They may also struggle to repeat phrases accurately, often omitting words or stumbling.
- **svPPA**: Semantic Variant, Primarily affects the understanding of word meanings (semantic knowledge). Patients may struggle with naming and comprehension, even for common objects. They often resort to broad categories instead of precise words (e.g., thing instead of fork). Example: When shown a picture of a dog, the patient might say, “It’s an animal… I think it’s a pet,” without being able to retrieve the word “dog.” They may also have difficulty understanding specific terms, relying on broader descriptions.
- **nfvPPA**: Impacts grammar and speech production, leading to slow, effortful, and agrammatic speech. Patients may omit small grammatical words (e.g., “is,” “the”) and speak in a telegraphic manner. Patients tend to use very short sentences, a rich vocabulary with low-frequency words, and more nouns compared to verbs. Example: The patient might say, “Walk… store… buy milk,” instead of “I’m going to walk to the store to buy milk.” Speech is often halting and labor-intensive, with noticeable pauses.
- **control**: The individual demonstrates fluent, grammatically correct speech, free from any markers of hesitation, effortful speech, or semantic impairment. There are no indications of word-finding difficulties or grammatical errors. The individual uses both simple and complex sentences naturally and appropriately. They can express themselves clearly without notable pauses, hesitations, or substitutions. The vocabulary used is appropriate for the context, and their language comprehension and responses are cohesive. Example: “I’m going to walk to the store to buy some milk” or “After I finish work, I plan to go for a walk and then cook dinner.” The language is fluent, natural, and demonstrates coherent sentence-building abilities.

Analyze the language, sentence structure, vocabulary, and speech flow within the conversational context of the interview to determine the most fitting category. Your response should include only one of the following labels: **lvPPA, svPPA, nfvPPA**, or **control**. If the text does not clearly fit into one category, analyze it carefully and suggest the most likely category based on available evidence.

### B Computational Resources

The experiments described in Section 4.2 and 6.2 were conducted on Google Colab Pro using an NVIDIA L4 GPU.

The experiments described in Section 5 were conducted using two different computational setups. For LLAMA and Mistral, we ran experiments locally on a system running Ubuntu 22.04.4 LTS (Jammy Jellyfish). This system featured an AMD Ryzen 9 7950X 16-Core Processor (32 threads, 16 cores, 2 threads per core) with a maximum clock speed of 5.88 GHz, 62 GB of RAM, 2 GB of swap space, and an NVIDIA RTX A6000 GPU with 48 GB of memory, using CUDA 12.4 for GPU acceleration. For GPT-3.5 and GPT-4o-mini, we relied on the OpenAI API, accessing the models via cloud-based inference.

### C Results of Traditional Machine Learning’s experiments

### D Results of LLMs’ Experiments

## Footnotes

1 https://dementia.talkbank.org/

2 https://huggingface.co/docs/transformers/main_classes/text_generation#parameters

